# Dzip1 is dynamically expressed in the vertebrate germline and regulates the development of *Xenopus* primordial germ cells

**DOI:** 10.1101/2024.04.20.590349

**Authors:** Aurora Turgeon, Jia Fu, Divyanshi, Meng Ma, Zhigang Jin, Hyojeong Hwang, Meining Li, Huanyu Qiao, Wenyan Mei, Jing Yang

**Affiliations:** Department of Comparative Biosciences, University of Illinois at Urbana-Champaign, Urbana, IL, USA; Chemistry Department of Comparative, Lake Washington Institute of Technology; Department of Cell and Developmental Biology, University of Illinois at Urbana-Champaign, Urbana, IL, USA; College of Life Sciences, Zhejiang Normal University, 688 Yingbin Road, Jinhua, Zhejiang 321004, China; Department of Biochemistry and Biophysics, University of Pennsylvania Perelman School of Medicine, Philadelphia, PA, USA; Department of Biochemistry & Molecular Biology, Shanxi Medical University, Taiyuan 030001, China

**Keywords:** Dzip1, Dazl, germ plasm, primordial germ cell, germline development, *Xenopus*

## Abstract

Primordial germ cells (PGCs) are the precursors of sperms and oocytes. Proper development of PGCs is crucial for the survival of the species. In many organisms, factors responsible for PGC development are synthesized during early oogenesis and assembled into the germ plasm. During early embryonic development, germ plasm is inherited by a few cells, leading to the formation of PGCs. While germline development has been extensively studied, how components of the germ plasm regulate PGC development is not fully understood. Here, we report that Dzip1 is dynamically expressed in vertebrate germline and is a novel component of the germ plasm in *Xenopus* and zebrafish. Knockdown of Dzip1 impairs PGC development in *Xenopus* embryos. At the molecular level, Dzip1 physically interacts with Dazl, an evolutionarily conserved RNA-binding protein that plays a multifaced role during germline development. We further showed that the sequence between amino acid residues 282 and 550 of Dzip1 is responsible for binding to Dazl. Disruption of the binding between Dzip1 and Dazl leads to defective PGC development. Taken together, our results presented here demonstrate that Dzip1 is dynamically expressed in the vertebrate germline and plays a novel function during *Xenopus* PGC development.

## Introduction

Germline development is crucial for the survival of the species. Successful germline development relies on the formation of functional gametes, which are derived from the primordial germ cells (PGCs) in early embryos. In different organisms, specification of the PGC fate can occur through two different mechanisms that have been broadly characterized as inheritance or induction (1-4). In mammals, BMP signaling specifies a pool of pluripotent cells in the epiblast as the PGCs during gastrulation (5-9). In contrast, animals such as *Drosophila* (10,11), *C. elegans* (11,12), *Xenopus*, and zebrafish (13-16) specify their PGCs through the inheritance of maternally supplied germ plasm, which contains a unique set of proteins and RNAs that are sufficient and necessary for germline determination (17-19). Although PGC specification may occur through different mechanisms in different species, the majority of factors responsible for PGC development are evolutionarily conserved (4,20).

In zebrafish and *Xenopus*, germ plasm is assembled during early oogenesis within a membrane-less structure called the Balbiani body (Bb) (4,15,21,22). Subsequently, the Bb is fragmented into numerous small germ plasm aggregates. These germ plasm aggregates are transported vegetally and ultimately reach the cortex of the vegetal pole, located between the cortical granules and yolk. During the oocyte-to-embryo transition, many germ plasm aggregates are degraded (23,24). The remaining ones coalesce into larger aggregates and are asymmetrically segregated into a small number of cells, which adapt to the PGC cell fate (25). PGCs then proliferate and migrate toward the gonads. In *Xenopus*, PGCs undergo approximately three rounds of proliferation during the migration, which are vital to ensure that enough PGCs can reach gonads and differentiate into gametes (26). Although it is known that maternally provided factors in the germ plasm play important roles in controlling many aspects of PGC development, the biological functions and regulatory mechanisms of many germ plasm components are not fully understood yet.

One of the germline determinants in the germ plasm is deleted in azoospermia-like (Dazl), an RBP essential for germline development, evolutionarily conserved from worms to mammals (27). During *Xenopus* PGC development, *dazl* mRNA and Dazl protein are localized in the germ plasm (28-30) and play a key role in PGC migration and proliferation (28). During zebrafish germline development, *dazl* is localized in the germ plasm (31). By characterizing the *dazl* mutants, Bertho et al. reported that Dazl is required for germline cyst formation and plays crucial roles in germ cell amplification and establishment of germline stem cells in zebrafish (32). In mice, Dazl regulates meiosis in both male and female germlines (33-36). Interestingly, Dazl is involved in the proper formation of the meiotic spindle during the oocyte-to-embryo transition (37). Overexpression of Dazl and Boule, another member of the Daz protein family (38), could modulate human embryonic stem cells (hESCs) to exit pluripotency and enter into meiosis (39).

DAZ-interacting protein 1 (Dzip1) is a component of the appendages of the mother centrioles and the cilia basal body that is essential for ciliogenesis (40-43). Cilia are required for vertebrate Hedgehog (Hh) signaling, which plays a key role in the patterning of the neural tube and the paraxial mesoderm. Loss of Dzip1 causes midline defects during vertebrate embryonic development (43-46). Dzip1 plays a multifaced role in the Hh pathway. In addition to its role in receiving Hh signals, Dzip1 can regulate the Hh pathway at the level of Gli transcriptional factors. It has been reported that Dzip1 can physically bind Gli3 and prevent Gli3 from translocating into the nucleus (43). Dzip1 also regulates the stability of Spop, an E3 ubiquitin ligase that promotes proteasome-dependent turnover of Gli proteins (46). Interestingly, while Dzip1 was originally cloned for its ability to interact with Daz (Deleted in Azoospermia) (47), its function during germline development has not been studied. Nevertheless, it has been reported that in humans, Dzip1 deficiency causes sperm centriole dysfunction and impairs the formation of sperm flagella, leading to asthenoteratospermia (48). Here, we report that Dzip1 is dynamically expressed in vertebrate germ cells and is required for the development of *Xenopus* PGCs.

## Results

### Expression of Dzip1 protein during vertebrate germline development

To assess the expression of Dzip1 protein during *Xenopus* development, we generated an anti-Dzip1 antibody, using a synthesized peptide (ESFEVPRSVKSRPLAK). On western blot, this antibody recognizes a 95kDa endogenous protein. The level of this protein was reduced upon injection of a Dzip1 morpholino (DMO) into *Xenopus* embryos (46,49) (Fig 1A). As expected, this antibody recognizes cilia basal bodies in the epidermal multi-ciliated cells in *Xenopus* embryos at stage 25 (Fig 1B). The immunofluorescence signal was markedly reduced in DMO-injected embryos (Fig 1C), further demonstrating the specificity of the antibody. Using this antibody, we assessed the expression of Dzip1 during *Xenopus* development. During oogenesis, Dzip1 was detected in the Balbiani body (Bb) (Fig 2A), a structure in the stage I oocytes essential for oocyte polarity and germline development (50). In stage II oocytes (Fig 2B), stage VI oocytes (Fig 2C), and mature eggs (Fig 2D), Dzip1 localizes to the vegetal cortex, in a manner reminiscent of the germ plasm. The staining signal in the vegetal cortex was reduced upon knockdown of endogenous Dzip1 (Fig 2E), providing additional confirmation for the specificity of the antibody. To determine if Dzip1 and germ plasm colocalize, we assessed Dzip1 expression using embryos derived from Dria transgenic frogs (51). We found that Dzip1 colocalizes with germ plasm in Dria embryos at all stages analyzed (Fig 2F, F’, F”, G, G’, G”, H, H’, H”). These observations demonstrate that the Dzip1 protein is a component of *Xenopus* germ plasm.

**Fig 1.**
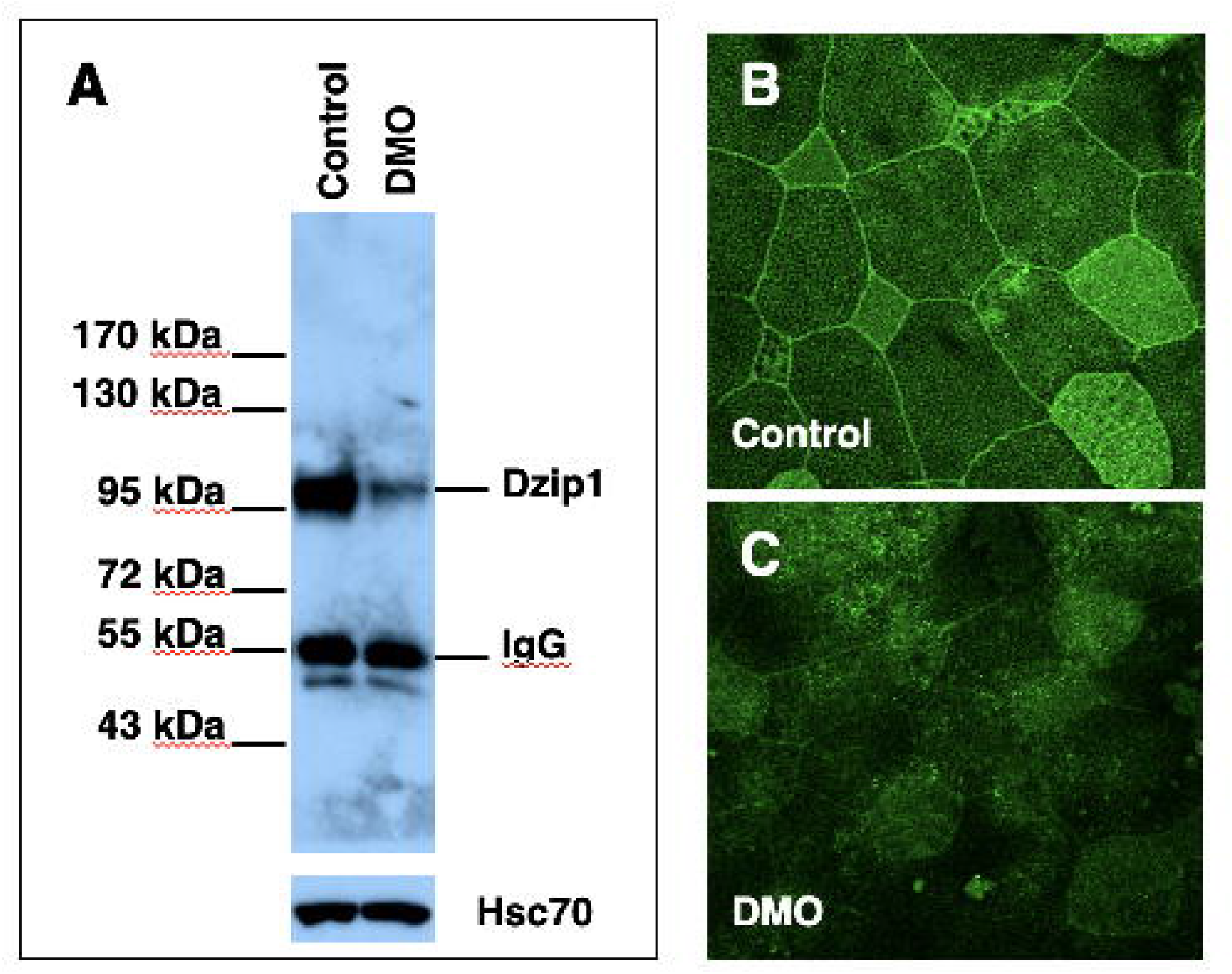
Specificity of the anti-Dzip1 antibody. **A**. Lysates from control and DMO (10 ng) injected embryos (stage 20) were immunoprecipitated using the affinity-purified anti-Dzip1 antibody. IP samples were subjected to western blot. **B**. Whole mount immunostaining of the control and DMO (10 ng) injected embryos (stage 25) using the affinity-purified anti-Dzip1 antibody.

**Fig 2.**
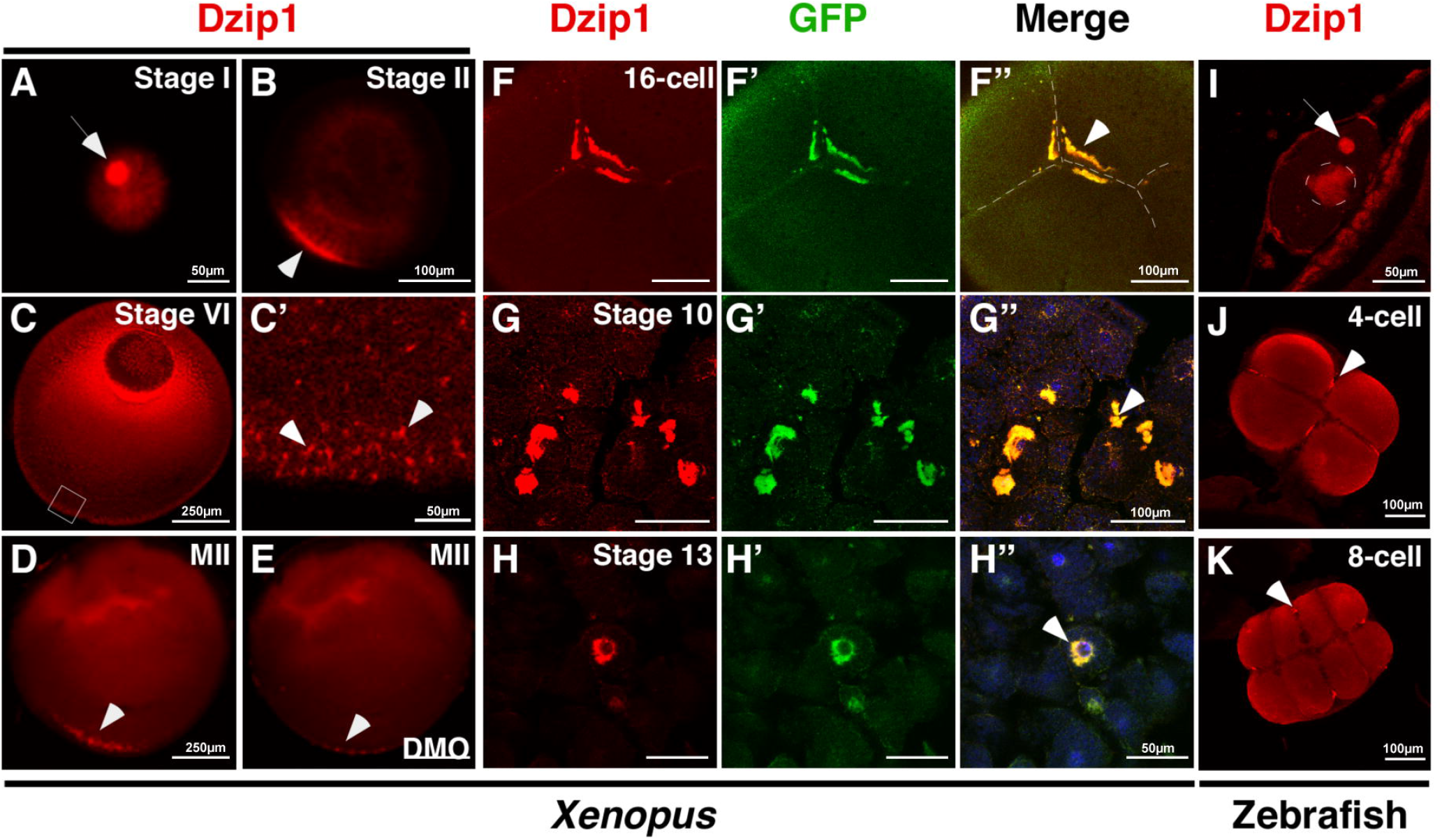
Expression of Dzip1 protein in *Xenopus* and zebrafish germline. **A - H** are *Xenopus* oocytes (**A** - **E**) and embryos (**F** - **H**). **I - K** are zebrafish oocytes (**I**) and embryos (**J** and **K**). **F** - **H** are 16-cell stage (**F**), stage 10 (**G**), and stage 13 (**H**) embryos from the Dria transgenic frog, which carries a mitochondria-specific GFP transgene. **I - K** are zebrafish oocyte (**I**), 4-cell stage (**J**), and 8-cell stage (**K**) embryos. Arrows in **A** and **I** point to the Bb. Arrowheads point to germ plasm.

Inspired by the above findings, we investigated the expression of Dzip1 in the germline of other vertebrate species. We first assessed the expression of Dzip1 in zebrafish. Indeed, Dzip1 is enriched in the Bb in zebrafish stage I oocytes (Fig 2I). After fertilization, while the Dzip1 protein is present in the entire blastodisc, highly concentrated Dzip1 protein aggregates were detected at four corners of the cleavage furrow (Fig 2J and K), where germ plasm aggregates are located during cleavage stages. This pattern is strikingly similar to the expression of Dzip1 during *Xenopus* germline development.

We further examined the expression of Dzip1 in mouse germline. In the ovary, Dzip1 is expressed in oocytes and surrounding somatic cells. Within the oocyte, Dzip1 can be detected in both the ooplasm and GV (Fig 3). While Dzip1 is relatively evenly distributed in the ooplasm throughout oogenesis, its expression is rather dynamic inside the GV. In the primary follicle, Dzip1 is strongly expressed inside the GV (Fig 3A-A”). In the secondary follicle, Dzip1 is highly enriched in a large spherical structure, likely the nucleolus (Fig 3B-B”). In the antral follicle, the size of the GV increases further. Except for some fibrous structures inside the GV, Dzip1 was not detected in the nucleoplasm (Fig 3C-C”).

**Fig 3.**
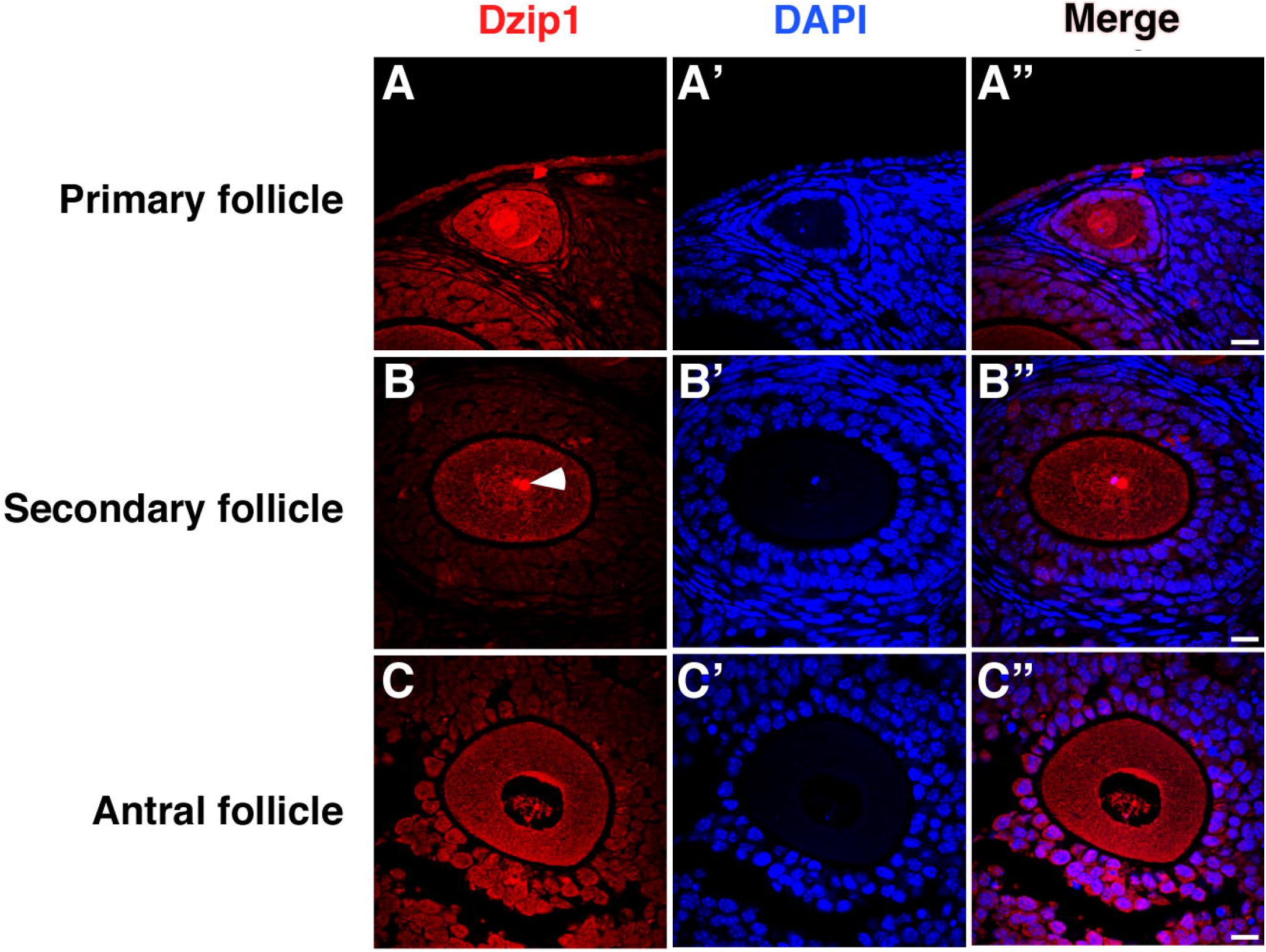
Expression of Dzip1 protein in the mouse ovary. **A** shows a primary follicle. **B** shows a secondary follicle. **C** shows an antral follicle. The arrowhead in B points to the nucleolus-like structure in the GV. DAPI stains nuclei on each section. The scale bar represents 20 μm.

In the testis, Dzip1 exhibits a much more dynamic expression pattern. Specifically, Dzip1 was detected in a subset of cells close to the basal lamina in the seminiferous tubules. These Dzip1-positive cells do not express the spermatogonia marker Plzf (Fig 4A-A’”), but express Sycp3 (Fig 4B-C’”), a component of the synaptonemal complex that is expressed in spermatocytes.

**Fig 4.**
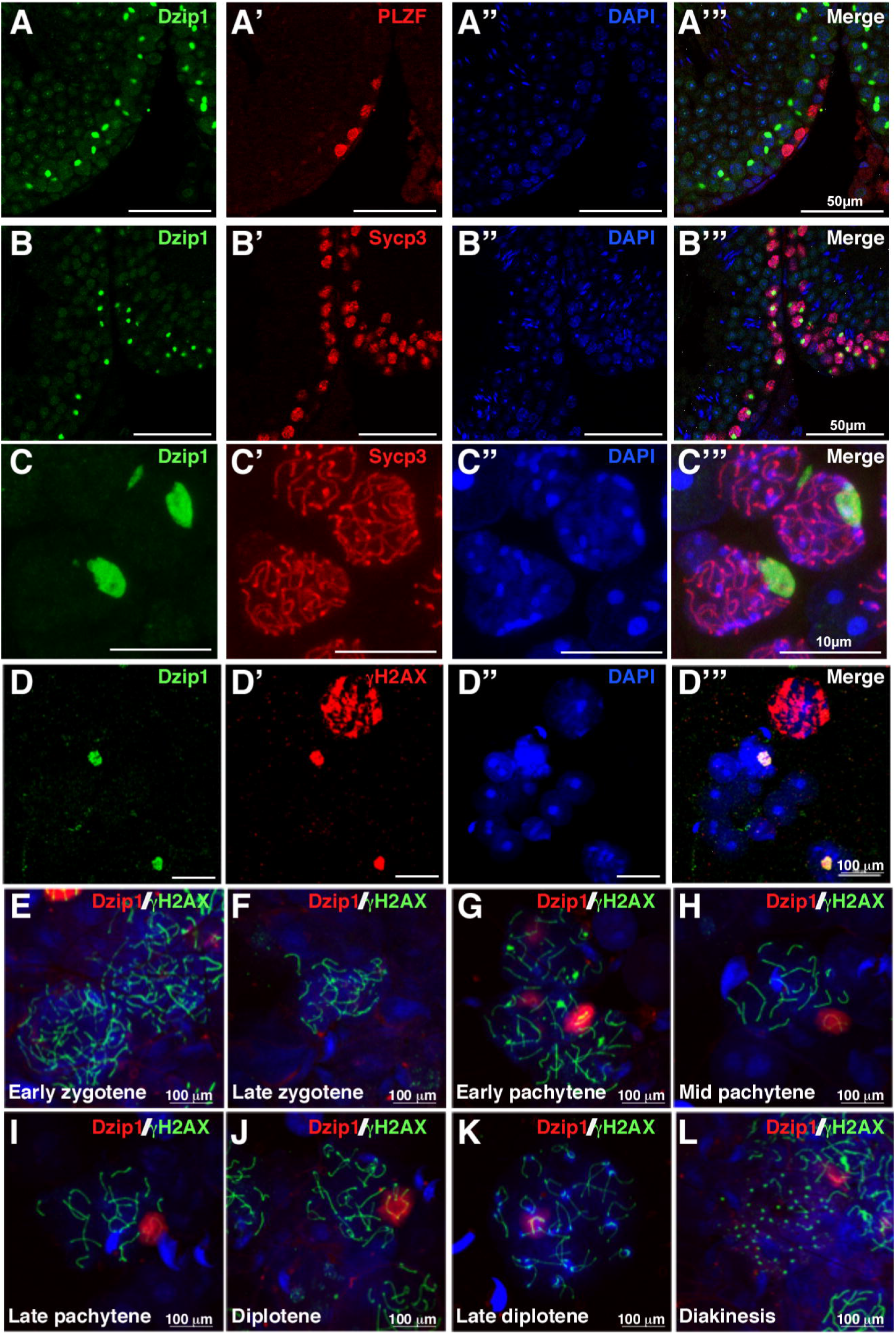
Expression of Dzip1 in mouse testis. **A - C** are immunofluorescence, showing the expression of Dzip1 in mouse testis tissue. **A**. Dzip1 (green) and PLZF (red) double staining. **B**. and **C**. Dzip1 (green) and Sycp3 (red) double staining. **C** is the high magnification view, which shows the spermatocyte nuclei in **B. D - L** are immunostained spreads of spermatocyte chromosomes from adult mouse testes. **D, D’, D”**, and **D’”** are Dzip1 (green) and γH2AX (red) double staining. **E** - **L** are Dzip1 (red) and γH2AX (green) double staining.

This demonstrates that Dzip1 is specifically expressed in spermatocytes during spermatogenesis. On high magnification images, it appears that the Dzip1 protein forms a large spherical aggregate inside the nucleus of the spermatocyte (Fig 4C-C’”). Double immunofluorescence staining on spermatocyte chromosomes spreads further reveals that the Dzip1 protein colocalizes with γH2AX (Fig 4D-D’”), a marker of the sex body, which contains silenced sex chromosomes. Of note, Dzip1 colocalizes with γH2AX in a very narrow time window during meiosis. Dzip1 was not detected during the zygotene stage (Fig 4E and F). Strong expression of Dzip1 was observed from pachytene to diplotene stages (Fig 4G, H, I, and J). Starting from the late diplotene stage, Dzip1 expression declines sharply (Fig 4K and L). This dynamic expression pattern suggests that Dzip1 may play a novel role in the formation of the sex-body during spermatogenesis. As Dzip1 is expressed in the germline in zebrafish, *Xenopus*, and mice, we speculate that Dzip1 is a novel regulator of vertebrate germline development.

### Dzip1 is required for Xenopus PGC development

To determine if Dzip1 is required for *Xenopus* germline development, we knocked down Dzip1 by injecting Dzip1 morpholinos (DMO1 and DMO2), which block translation (49). At stage 33, we harvested embryos and assessed the number of PGCs by *in situ* hybridization for *pgat* (52). As shown in Fig 5A and B, embryos injected with DMO1 or DMO2 exhibited a significant reduction in the number of PGCs. This phenotype was rescued by vegetal injection of 500pg of RNA encoding myc-Dzip1 (Fig 5C and D), demonstrating the specificity of knockdown. Dzip1 is involved in ciliogenesis and Hh signaling (40-42,44-46,49). To determine if Dzip1 regulates PGC development by controlling ciliogenesis or Hh signaling, we knocked down IFT88, which is essential for ciliogenesis (53,54), and Gli1, a key downstream mediator of Hh signaling (55). We found the number of PGCs remained normal in IFT88- and Gli1-knockdown embryos (Fig 5E), indicating that Dzip1 regulates PGC development independent of its roles in controlling ciliogenesis and Hh signaling. To better understand how Dzip1 regulates PGC development, we harvested Dzip1 knockdown embryos at multiple time points. Strikingly, the number of *pgat*-positive PGCs was normal at stage 16 in Dzip1 knockdown embryos. At stage 22, the number of PGCs starts to increase in control embryos. Compared to control embryos, Dzip1 knockdown embryos show slightly less number of PGCs. At stage 28, as the PGC number further increased in control embryos, the reduction in the number of PGCs is more prominent in Dzip1 knockdown embryos (Fig 5F). These results demonstrate that the knockdown of Dzip1 does not affect PGC specification but impairs PGC development during tailbud stages.

**Fig 5.**
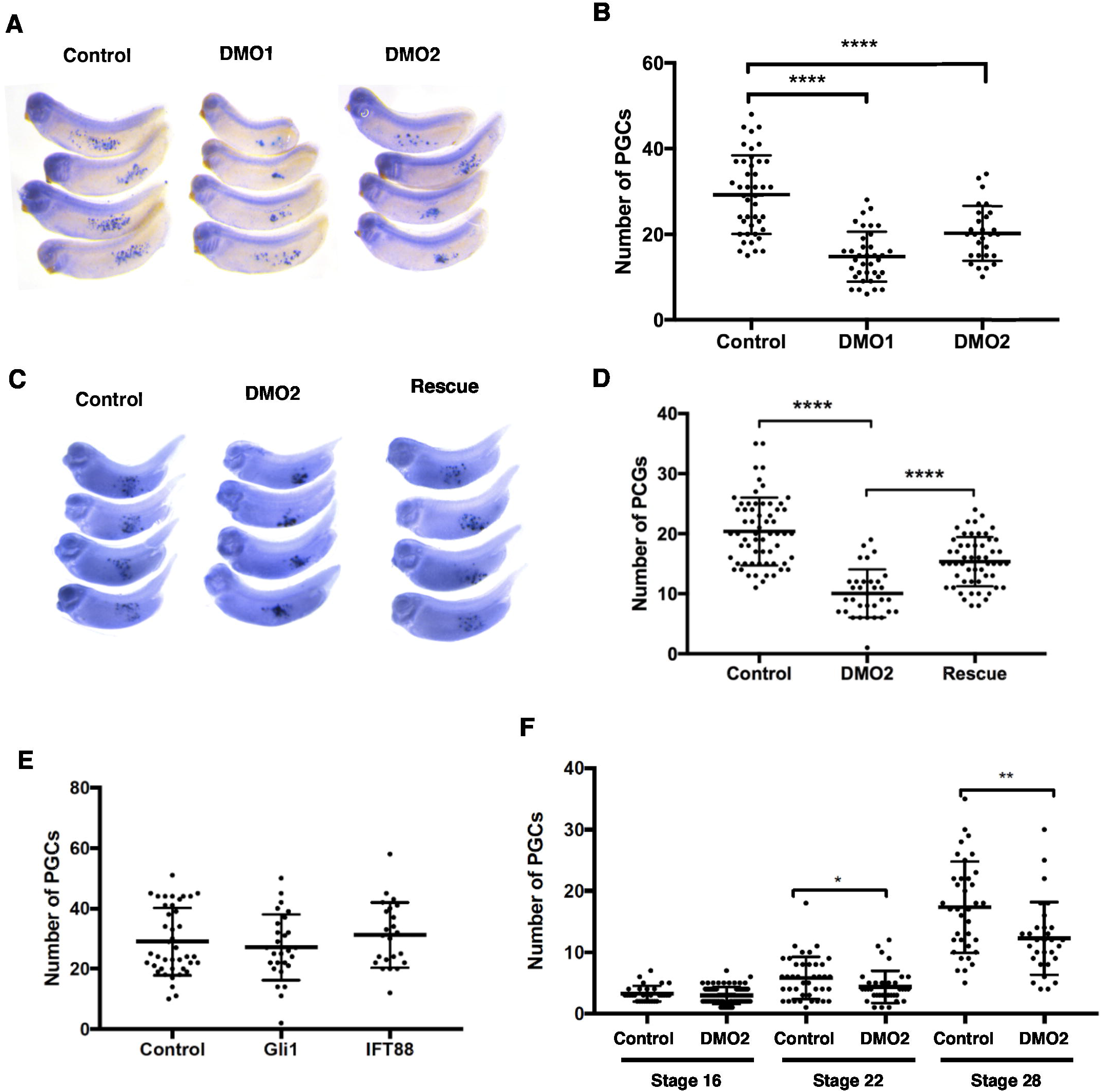
Dzip1 is required for PGC development in *Xenopus*. **A**. knockdown of Dzip1 impaired *Xenopus* PGC development. Whole-mount *in situ* hybridization shows the expression of *pgat* in control and Dzip1 morpholino (DMO1 and DMO2, 10 ng per embryo) injected embryos. **B**. Quantification of the results shown in A. The number of *pgat*-positive PGCs in each embryo was counted and plotted on the graph. **C**. The specificity of Dzip1 knockdown. Whole-mount *in situ* hybridization shows the expression of *pgat* in control, DMO2 (10 ng) injected, and DMO2 (10 ng) + myc-Dzip1 (500 pg) injected embryos. **D**. Quantification of the number of *pgat*-positive PGCs shown in **C. E**. Knockdown of Gli1 and IFT88, which impaired Hh signaling and ciliogenesis respectively, had no detectable effect on *Xenopus* PGC development. Quantification of the number of *pgat*-positive PGCs in control, Gli1 morpholino (10 ng) injected, and IFT88 morpholino (10 ng) injected embryos at stage 33. **F**. Knockdown of Dzip1 does not affect *Xenopus* PGC specification. Quantification of the number of *pgat*-positive PGCs in control and DMO2 (10 ng) injected embryos at stages 16, 22, and 28. Statistical analysis in **B, D, E**, and **F** was done by two-tailed *t*-tests. \**P*<0.05; \*\**P*<0.01; \*\*\*\**P*<0.0001. Data are mean±s.d.

### Physical and functional interaction between Dzip1 and xDazl

Dzip1 interacts with Daz (47). To understand the mechanism by which Dzip1 regulates germline development, we first characterized the biochemical interaction between Dzip1 and xDazl, the *Xenopus* homolog of Daz.

We transfected HEK293T cells with Flag-tagged xDazl (Flag-xDazl) and myc-tagged Dzip1 (Myc-mDzip) either individually or in combination. In a co-immunoprecipitation experiment, we found Flag-xDazl was co-purified with Myc-Dzip1 (Fig 6A, upper panel). Thus, Dzip1 can form a complex with xDazl (Fig 6A, lower panel). We further mapped the xDazl binding domain in Dzip1 using several Dzip1 deletion constructs, including 1-550, 550-850, 1-221, and 282-550 (Fig 6B). Results from our co-immunoprecipitation indicate that the sequence between amino acid residues 282 and 550 is responsible for binding xDazl (Fig 6C).

**Fig 6.**
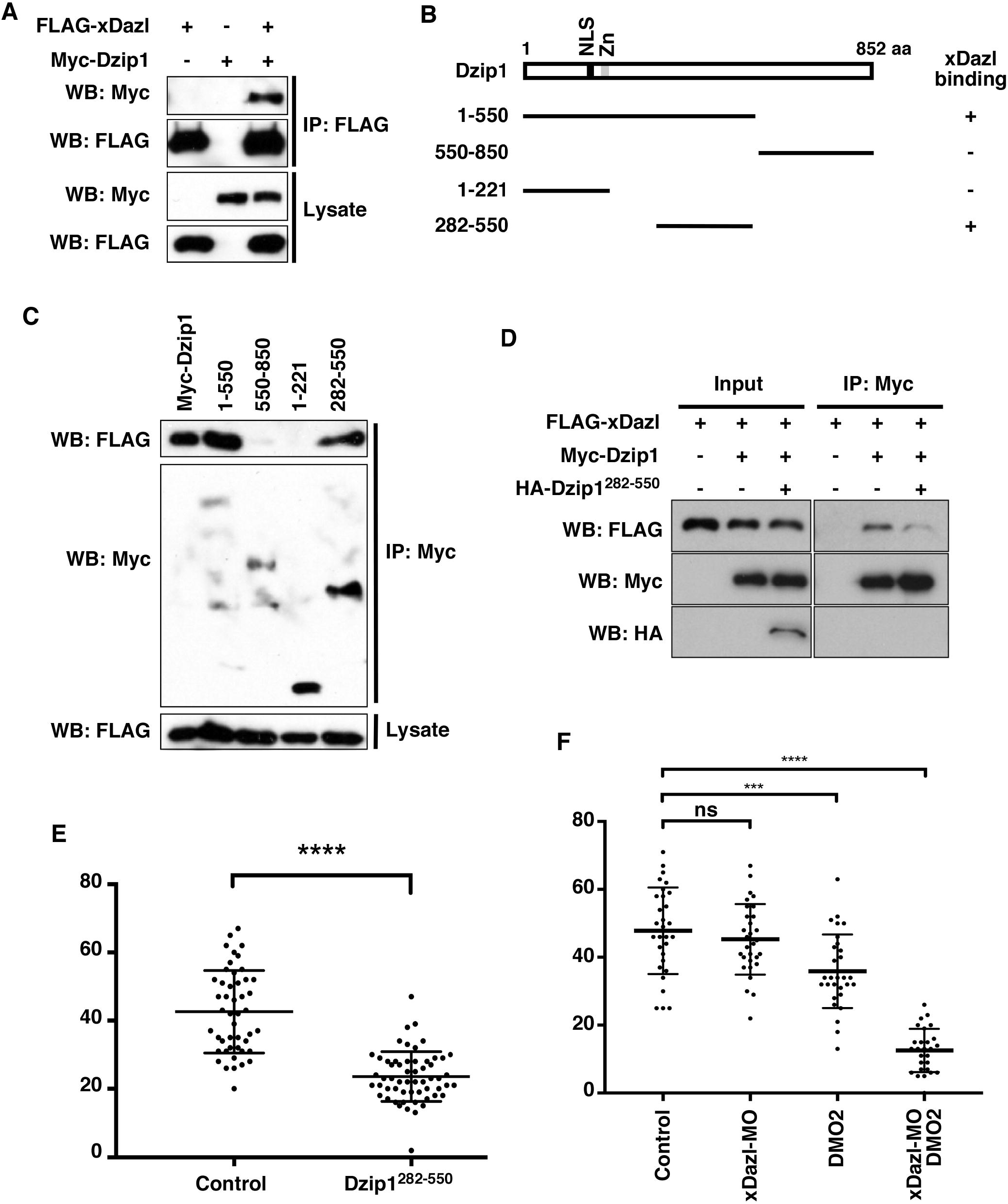
The interaction between Dzip1 and xDazl is required for *Xenopus* PGC development. **A**. Dzip1 physically interacts with xDazl. FLAG-xDazl and Myc-Dzip1 were expressed in HEK293T cells for CoIP analysis. **B**. Schematic drawing to show the structure of Dzip1 protein and Dzip1 deletion constructs. Whether Dzip1 constructs bind xDazl was indicated by “+” (bind) or “-” (not bind). **C**. Co-IP results show that xDazl interacts with the full-length Dzip1, 1-550, and 282-550, but not 1-221 and 550-850. **D**. Co-IP results show overexpression of Dzip1^282-550^ interfered with the complex formation between the full-length Dzip1 and xDazl. **E**. Interfering with the complex formation between Dzip1 and xDazl impaired PGC development in *Xenopus* embryos. Control and Dzip1^282-550^ (1 ng) injected embryos were harvested at stage 33 for *pgat in situ* hybridization. The number of *pgat*-positive PGCs in control and Dzip1^282-550^ injected embryos were counted and plotted on the graph. **F**. Knockdown of Dzip1 and xDazl impaired *Xenopus* PGC development synergistically. Embryos were injected with DMO2 (2.5 ng) and xDazl morpholino (xDazl-MO, 3 ng) alone, or in combination. Control and injected embryos were harvested at stage 33 for *pgat in situ* hybridization. The number of *pgat*-positive PGCs was counted and plotted on the graph. Statistical analysis in **E** and **F** was done by two-tailed *t*-tests. \**P*<0.05; \*\*\**P*<0.001; \*\*\*\**P*<0.0001. ns – not significant. Data are mean±s.d.

Knockdown of xDazl reduced the number of PGCs in *Xenopus* embryos and impaired PGC migration (28). Since Dzip1 physically interacts with xDazl and is required for PGC development, we set out to determine if the interaction between Dzip1 and xDazl is important for PGC development. We reasoned that Dzip^282-550^, the xDazl-binding domain of Dzip1, might act as a dominant negative Dzip1 by preventing the complex formation between the full-length Dzip1 and xDazl. As expected, when Dzip^282-550^ was coexpressed with Dzip1 and xDazl in HEK293T cells, the interaction between Dzip1 and xDazl was reduced (Fig 6D). We injected Dzip^282-550^ into the vegetal pole of 1-cell stage *Xenopus* embryos and assessed the number of PGCs at the tailbud stage. Indeed, we found that overexpression of Dzip^282-550^ significantly reduced the number of PGCs in the embryos (Fig 6E).

Parallel to the above analysis, we determined if partial knockdown of Dzip1 and xDazl could reduce the number of PGCs synergistically. We chose low doses of Dzip1 morpholino (DMO2, 2.5 ng) and xDazl morpholino (Dazl-MO, 3 ng), which alone only reduced the number of PGCs slightly. When both morpholinos were co-injected into embryos, we found the number of PGCs was severely reduced (Fig 6D). This result further supports the idea that the interaction between Dzip1 and xDazl is important for PGC development.

## Discussion

Dzip1 is a component of the cilia basal body. It is required for ciliogenesis and Hh signaling (40-46,49). Although it was originally identified for its ability to interact with Daz (47), which is important for germline development, the expression and function of Dzip1 during germline development have not been studied. It remains unclear if the interaction between Dzip1 and the Daz family of proteins is important for germline development.

Our work demonstrates that Dzip1 is dynamically expressed during *Xenopus*, zebrafish, and mouse germline development. Dzip1 can be detected in the nucleus and ooplasm in the oocyte in all three vertebrate species. In *Xenopus* and zebrafish oocytes, Dzip1 is enriched in the Bb during oogenesis. During early embryonic development, Dzip1 localizes to the germ plasm at all stages analyzed. Consistent with its expression pattern, we found that Dzip1 is required for PGC development in *Xenopus*.

Our results further reveal that Dzip1 regulates *Xenopus* PGC development via its interaction with xDazl, independent of its functions in controlling Hh signaling and ciliogenesis. In agreement with this view, we show that the knockdown of Dzip1 did not affect PGC development during the early stages. Starting from the tailbud stage, the number of PGCs was reduced in Dzip1-depleted embryos. This phenotype is striking similar to what was observed in xDazl deficient embryos (28). We were able to verify the binding between Dzip1 and xDazl, and mapped the xDazl-binding domain to the sequence between residues 282 and 550 of Dzip1. Overexpression of Dzip1^282-550^, which competes with the full-length Dzip1 for binding xDazl, interfered with PGC development in *Xenopus* embryos. Moreover, partial knockdown of Dzip1 and xDazl impaired PGC development synergistically. These findings demonstrate that Dzip1, a component of the germ plasm, is a novel regulator of germline development.

Our results presented here reveal that the physical interaction between Dzip1 and xDazl is important for PGC development. We speculate that by forming a complex with xDazl, Dzip1 may influence the function of xDazl. As an RNA-binding protein, Dazl regulates the translation of its target RNAs in a context-dependent manner. A recent study has demonstrated that during mouse oocyte maturation, Dazl plays both positive and negative roles in the translation of its targets. Whether Dazl promotes or suppresses translation depends on the context of the 3’UTR of its target RNA (56). Dazl can interact with translational repressors such as Pumilio2 (57), which represses protein expression by inhibiting translation and promoting mRNA decay (58). On the other hand, Dazl can activate translationally silent mRNAs through the direct recruitment of poly(A)-binding proteins (PABPs), which are critical for the initiation of translation (59). Future work is needed to determine if Dzip1 regulates the binding between Dazl and translational activators and repressors.

It is worth mentioning that during mouse male germline development, Dzip1 is expressed specifically in the sex body between the pachytene and diplotene stages. Both germ plasm in zebrafish and *Xenopus* and sex body in mice are phase-separated insoluble protein condensates (60-62). Detection of Dzip1 in germ plasm and sex body could suggest that Dzip1 may play a role in the formation or maintenance of phase-separated protein condensates in the germline. In the future, it will be of great interest to determine if Dzip1 is involved in phase separation during germline development.

## Supporting information

Key Resource Table

## Acknowledgment

This work is supported by a grant from NIH (R35 GM131810).

## Materials and Methods

### Animals

*Xenopus laevis* oocytes and embryos were obtained, and injected as described [85]. Briefly, embryos were placed in 0.5× Marc’s Modified Ringer’s with 3% Ficoll. Morpholinos or RNAs were injected into the vegetal pole using a Narishige IM300 microinjector. For experiments in which RNA and morpholino were injected into the same embryo, injections were performed sequentially. The dosages of morpholinos and RNAs for microinjection are described in the text or figure legends. Morpholinos against Dzip1 (DMO1 and DMO2) (46,49), Gli1 (55), and intraflagellar transport protein 88 (IFT88)/Polaris (54) were described. The sequence of xdazl morpholino is TTTCCAGACATTCTTTCAACGATGA. *Xenopus* procedures were approved by the University of Illinois at Urbana-Champaign Institutional Animal Care and Use Committee (IACUC) under animal protocol #23088 and performed in accordance with the recommendations of the Guide for the Care and Use of Laboratory Animals of the National Institutes of Health.

Zebrafish procedures were approved by the University of Illinois at Urbana-Champaign Institutional Animal Care and Use Committee (IACUC) under animal protocol #22160 and performed in accordance with the recommendations of the Guide for the Care and Use of Laboratory Animals of the National Institutes of Health. The female Tübingen strain was purchased from ZIRC (Zebrafish International Resource Center). Ovary tissue was fixed in 4% PFA and embedded in paraffin for sectioning. Embryos were fixed in 4% PFA for immunostaining.

Mice procedures were approved by the University of Illinois at Urbana-Champaign Institutional Animal Care and Use Committee (IACUC) under animal protocol #23096 and performed in accordance with the recommendations of the Guide for the Care and Use of Laboratory Animals of the National Institutes of Health. Mouse testis was harvested and stained as described (63).

### Cell culture

HEK293T cells were cultured in DMEM, supplemented with 10% FBS and 1× Penicillin-Streptomycin solution (complete medium). Cell cultures were maintained in a standard humidified incubator at 37 °C with 5% CO_2_. Transfection mixtures were prepared by mixing various combinations of expression constructs (total 2 µg) within 200 µL DMEM containing 1.2 µg of polyethyleneimine (PEI, PEI-to-DNA mass ratio of 1.5:1). The transfection mixtures were incubated at room temperature for 20 minutes before adding to cells cultured in a 6-well plate with 2 mL complete medium. The transfection medium was replaced with 2 mL complete medium after 24 hours of transfection.

### Antibodies

Antibodies used were anti-Myc (9E10, Thermo Fisher, 1:1000), anti-FLAG (M2, Sigma, 1:1000), anti-HA (Thermo Fisher, 1:1000), and rabbit anti-mDzip1 (49). A rabbit anti-xDzip1 antibody was raised using a peptide (Ac-CESFEVPRSVKSRPLAK-amide) that is close to the C-terminus of the protein. Anti-xDzip1 antibody was affinity purified using the peptide.

### CoIP, western blots, immunofluorescence, and whole mount *in situ* hybridization

CoIP and western blot were performed as described (49). Lysates were made using the NP-40 lysis buffer (50 mM Tris pH7.6, 125 mM NaCI, 1 mM EDTA, 0.1% NP-40). For whole mount immunofluorescence, *Xenopus* oocytes, mature eggs, embryos, and zebrafish embryos were fixed with Dent’s fixative (80% methanol, 20% DMSO) overnight, followed by incubation in Methanol at -20°C for at least 24 hours. Oocytes and mature eggs were rehydrated in 1 x TBS and blocked for 1 hour in the blocking buffer containing 10% donkey serum at room temperature, followed by incubation with primary antibodies at 4°C overnight. Oocytes and mature eggs were washed 5 times with 1 x TBSX (TBS with 0.1 % triton X-100) and incubated with the secondary antibodies overnight. After 5 washes with 1 x TBSX, oocytes and eggs were mounted with the BABB clearing solution (1 x benzyl alcohol: 2 x benzyl benzoate) for confocal imaging using NikonA1Rsi. To perform immunofluorescence staining of zebrafish ovary, ovary tissue was fixed in 4% PFA, paraffin-embedded, sectioned, and stained following the standard protocol. *Xenopus* whole mount *in situ* hybridization was performed as described (64). Plasmid for *pgat in situ* probe was described (52). For quantifications and statistical analysis, GraphPad Prism 7 was used.

**See other information in a separate KRT table**

