## Supplementary material for "Dzip1 is dynamically expressed in the vertebrate germline and regulates the development of *Xenopus* primordial germ cells": Key Resource Table

| Reagent or resource | Source | Identifier |
| --- | --- | --- |
| Antibodies | | |
| Mouse anti-c-Myc | Thermo Fisher | Cat. 13-2500; RRID: AB_2533008 |
| Mouse anti-FLAG M2 | Sigma | Cat. F1804;  RRID:AB_262044 |
| Mouse anti-HA | Thermo Fisher | Cat. 26183 AB_10978021 |
| Rabbit anti-mDZIP1 | Jin et al. 2011 (49) | N/A |
| Mouse anti-HSC70 | Santa Cruz | Cat. sc-7298; RRID: AB_627761 |
| Experimental Models: Cell Lines | | |
| HEK293T | Originally from ATCC | CRL-3216 |
| Experimental Models: Organisms/Strains | | |
| Xenopus laevis: Wild type | Nasco | LM00715 |
| Xenopus laevis: Dria transgenic line | Taguchi et al. 2012 (51) | NXR |
| *Danio rerio*: TÜ (Tübingen) lines | ZIRC (Zebrafish International Resource Center) | ZL57 |
| Oligonucleotides | | |
| DMO1 (DZIP1 morpholino 1) | Gene Tools, Philomath, OR | 5′-AGCAGCCTCTTCTCCTTCTGCGCTG-3′ |
| DMO2 (DZIP1 morpholino 2) | Gene Tools, Philomath, OR | 5′-AAAGGCATTCTCTTTCCTCCCGCTC-3′ |
| Xdazl morpholino | Gene Tools, Philomath, OR | 5’-TTTCCAGACATTCTTTCAACGATGA-3’ |
| Gli1morpholino | Gene Tools, Philomath, OR  Nguyen et al. 2005 (55) | 5’- CGGGCGGACACTGGCGGGACGC-3’ |
| IFT88/Polaris morpholino | Gene Tools, Philomath, OR  Dammermann et al. 2009 (54) | 5’-GCACGAGATGGACATTTTGCATCAT-3’ |
| Recombinant DNA | | |
| DZIP1 1-221 aa | Jin et al., 2011 (49) | N/A |
| DZIP1 282-550 aa | Jin et al., 2011 (49) | N/A |
| DZIP1 1-550 aa | Jin et al., 2011 (49) | N/A |
| DZIP1 550-850 aa | Jin et al., 2011 (49) | N/A |
| DZIP1 1-852 aa | Jin et al., 2011 (49) | N/A |
| Software and Algorithms | | |
| GraphPad Prism 7 | GraphPad | [<https://www.graphpad.com/>](https://www.graphpad.com/) |
